# Green synthesis of silver nanoparticles using psychrotolerant strains of *Tulasnella albida* from South Orkney Islands (Antarctica)

**DOI:** 10.1101/2021.09.02.458759

**Authors:** Jesica M. Kobashigawa, Carolina A. Robles, Rocío F. Gaiser, Daniel C. Schinca, Lucía B. Scaffardi, Cecilia C. Carmarán

## Abstract

Nanoparticles are widely studied due to their possible uses in biological and technological systems. Four psychrotolerant strains of *Tulasnella albida* isolated from Antarctica were tested and compared in their ability to synthesize silver nanoparticles. The four strains were capable of synthesizing silver nanoparticles with the addition of AgNO_3_ (final concentration of 0.5 mM), showing similar results under the same conditions: 28°C, 200 rpm, pH 9. Additionally, we registered the synthesis of nanoparticles at 6°C using biomass generated at the same temperature. For the characterization of synthesized nanoparticles, TEM and SEM microscopy analyses were performed. The images obtained showed the existence of spherically shaped silver nanoparticles with a log-normal size distribution centered at 2 nm diameter for 28°C. The largest ones showed a capping shell around them, appearing associated with the formation of small silver nanoparticles. Theoretical calculations of optical absorption based on core-shell *Ag*-*Ag*_*2*_*O* nanoparticles were used to characterize the experimental absorption spectra of silver nanoparticles colloids. This work contributes to expanding the studies and possible technological applications of psychrotolerant organisms in the industry, particularly in the green synthesis of nanoparticles at suboptimal conditions.

## INTRODUCTION

Nanoparticles (NPs) are a wide class of materials that have at least one dimension less than 100 nm, having physicochemical properties different from the same material but at bulk scale (Khan et al. 2019). These particular properties make them suitable for multiple applications such as medicine, electronics, manufacturing, among others (Khan et al. 2019). The synthesis of these nanomaterials through ecological, effective and low-cost methods has become relevant in recent years, in order to optimize resources and avoid contamination from hazardous chemical compounds or very high energy costs (Ahmad et al. 2019). Among the organisms used in the synthesis of nanoparticles, fungi are an interesting alternative due to their easy handling, tolerance to metals, their bioaccumulation capacity, their ability to secrete a very wide variety of enzymes, and their easy manipulation (Hulkoti et al. 2014). The potential of synthesis of these organisms has been reported in recent years (Rodrigues et al. 2013, Tyagi et al. 2019). These works, among others, have reported that changes in temperature, concentration of the metal precursor, pH, culture medium, and amount of biomass can be used to obtain nanoparticles with different physicochemical characteristics. These researches include the study of the synthesis of silver nanoparticles (AgNPs). These nanoparticles have a very wide range of applications as anti-bacterial agents in the health industry, food storage, textile coatings and a number of environmental applications as well as electronics and photocatalysis (Abou et al. 2010).

Extremophiles microorganisms are those that are able to grow and thrive in extreme environments, e.g. acidic or alkaline pH, high or low temperatures, high concentrations of pollutants, and salts, among others. Due to their unique physiological and enzymatic characteristics, which allow them to survive in hazardous environments, these organisms are promising for the synthesis of NPs (Purcarea et al. 2019). Among extremophiles, psychrophiles are those that have an optimum growth temperature below 15°C and cannot grow at temperatures beyond 20°C. On the other hand, psychrotolerant those which can live in cold conditions, but have a higher optimum growing temperature (Tiquia-Arashiro & Rodrigues 2016). The particular characteristics of the metabolism of these organisms represent valuable assets for advanced industrial applications (Purcarea et al. 2019). In the synthesis of NPs, extremophiles can have more pronounced ability to withstand conditions that may occur during the industrialization process and also, after the act of synthesis, adding more stability to the product with specific capping proteins that are less likely to leach when washed (Beeler & Singh 2016). The ability of the organisms and their proteins to survive in varied conditions also allows for some flexibility in the industrial environment; the organisms would survive and continue to produce NPs outside of optimal conditions, albeit with a lower yield (Beeler & Singh 2016). AbdelRahim et al. (2017), characterizing the synthesis of AgNPs in *Rhizopus stolonifer* at different temperatures, highlight the limitations of synthesis at low temperatures, not registering synthesis of AgNPs at 10 C due to denaturation or inactivation of enzymes and active molecules which are involved in the biogenesis of AgNPs. The fungi in low-temperature environments including icy habitats and deep-sea environments are of diverse nature and abundance. Their strategies to thrive under extreme conditions make them versatile and their metabolites can be of potential use in many dimensions (Rafiq et al. 2019).

Antarctica is the largest poorly explored pristine region on Earth and, in a similar way, its mycobiota remains mostly unknown, showing an inestimable value as a scientific preserve and biotechnological repository (Correa & Abreu 2020). Some fungal organisms can grow in the extreme conditions present in the white continent, facing the low temperatures that limit the available water and showing different morphological and physiological adaptive strategies (Ruisi et al. 2007). Among the organisms inhabiting Antarctica, the genus *Tulasnella* has been recently registered on decomposing wood from different constructions and artifacts from Deception Island (Held & Blanchette 2017) and on the historical heritage Museum Casa Moneta, located on the South Orkney Islands (Gaiser et al. 2020). *Tulasnella* comprises saprotrophic as well as mycorrhizal species, including endomycorrhizal partners of the Orchidaceae and also ectomycorrhizal fungi (Preussing et al. 2010). Limited reports about the enzymatic profiles of members of this genus are known. However, Kohler et al. (2015) showed that *Tulasnella* calospora has a robust apparatus for the degradation of crystalline cellulose, which includes cellobiohydrolases and lytic polysaccharide monooxygenases.

As part of a study that evaluates the synthesis of AgNPs using ligno-cellulolytic fungi (Kobashigawa et al. 2019), we explored the capacity of Antarctic psychrotolerant strains of *Tulasnella albida*, to reduce Ag+ ions into AgNPs. The aim of this work was to study the synthesis of AgNPs using *T. albida* strains (isolated from Antarctica) to different temperatures, by comparing their performance. Qualitative enzymatic tests of oxidases, tyrosinase and peroxidases were performed to determine if there could be a relation between the production of these enzymes with the synthesis of NPs. Additionally, characterization of the AgNPs obtained was performed.

Reports using psychrophilic (Apte et al. 2013) and psychrotolerant (Das et al. 2020) organisms were carried out previously. Nevertheless, to our knowledge no studies using psychrotolerant fungi from Antarctica evaluating the synthesis of NPs have been made.

## MATERIAL AND METHODS

### Chemical compounds

Silver nitrate (AgNO_3_, Sigma Aldrich) 100 mM was used for the biosynthesis of AgNPs. NaOH (1M) was used to adjust pH.

### Microorganisms

Four psychrotolerant strains of *Tulasnella albida* (BAFCcult 4710, 4711, 4712 and 4713, hereafter as strain 1, strain 2, strain 3 and strain 4, respectively) were used in all the assays.

These strains were isolated from wood from the Casa Moneta Museum located at the Orkney Permanent Base of the Laurie Island of the South Orkney archipelago in Antarctica, studied and characterized in the Mycology and Phytopathology Laboratories of the FCEN-UBA, and deposited in the culture collection of the Faculty of Exact and Natural Sciences, University of Buenos Aires (BAFCcult). Orkney station has an annual mean temperature of - 3.1 ± 0.7°C varying from 1.5 ± 0.5°C in summer to - 8.3 ± 2°C in winter (period 1981-2010) (Turner et al. 2020). The strains were incubated in 2% malt extract agar (MEA) plates, at 24°C.

### Growth curve of *Tulasnella albida*

Erlenmeyer flasks containing 50 mL of growth medium were used. Growth medium was composed of 10 g.L^-1^ glucose, 5 g.L^-1^ potato peptone, 3 g.L^-1^ malt extract and 3 g.L^-1^ yeast extract. The medium was sterilized, inoculated with two 5 mm plugs of the strain and incubated at 200 rpm at 28°C, in the dark, for 10 days. The mycelium was harvested every 2 days, filtered through a filter paper using a Büchner funnel and dried overnight at 40°C. Then, the dry weight of mycelia was determined. Three replicates were performed for each strain

### Biosynthesis of silver nanoparticles

The fungal biomass was obtained under the same conditions as for the growth curve, but the incubation only lasted 7 days. The mycelium obtained was filtered and washed with sterile water and re-incubated under the same conditions in 50 mL of sterile distilled water for 3 days. At the end of this incubation, the mycelium was filtered again, discarding the biomass and keeping the filtered liquid (hereafter, fungal filtrate).

For the biosynthesis of AgNPs, pH of 5 and 9 were evaluated. NaOH 1 M was used to adjust the pH. AgNO_3_ 100 mM was added to the fungal filtrate to obtain a final concentration of 0.5 mM. The fungal filtrate was incubated for 7 days under the same conditions as mentioned before. Aliquots containing only the fungal filtrate, without salt, and flasks containing sterile distilled water with the salt were established as controls. For all the treatments five replicates were performed.

To detect the biosynthesis of AgNPs, a 1 mL aliquot was removed from each flask and their UV-visible spectra were obtained using a spectrophotometer Shimadzu UV-MINI 1240 (Tokyo, Japan) by scanning the absorbance spectra in 200 – 800 nm range of wavelength. The synthesis of AgNPs was detected by the Surface Plasmon Resonance (SPR), which is a consequence of the oscillation of the electron cloud on the NP surface due to the radiation at a particular energy.

This resonance is due to their small size but it can be influenced by numerous factors that contribute to the exact frequency and intensity of the band, SPR peak of spherical AgNPs is conventionally observed at 400 nm (Jana et al. 2001) shifting the position of the plasmon peak to higher wavelengths as the particle radius increases (Slistan-Grijalva et al. 2005). Only data obtained at pH 9 is informed, as at pH 5 no positive results were obtained.

Given that as-obtained colloid samples may have different NPs size coexisting together, NPs synthesized by *T. albida* (strain 2) were centrifuged at 15000 rpm for 20 min as a first step to separate size components. Absorption spectrum of supernatant was then analyzed. As a second step, the supernatant previously obtained was re-centrifuged and a new absorption spectrum was recorded.

### Modeling

Optical absorption spectra of *Ag* colloid samples obtained in this work displaying SPR were modeled using Mie theory for small spherical NPs (Schinca & Scaffardi 2008). Since oxidation process cannot be avoided when AgNPs are synthesized in water, for our particular samples, core-shell *Ag-Ag*_*2*_*O* configuration was assumed. In the Rayleigh approximation for small NPs compared with wavelength, the polarizability of a core-shell particle with inner radius *r*_*1*_, outer radius *r*_*2*_, core dielectric function *ε*_1_ and shell dielectric function*ε*_2_ can be written as:

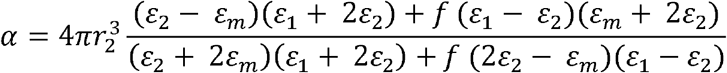

where *ε*_1_ is the surrounding medium dielectric function and 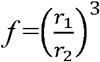.Metallic dielectric function *ε*_1_ was obtained from (Johnson & Christy 1927) while *ε*_2_ was taken from (Qiu et al. 2005).

### AgNPs synthesis efficiency

To study the formation of AgNPs for each strain, absorption spectra were taken on days 1, 2, 3 and 7 after the beginning of the synthesis. Efficiency was here defined as the height of the SPR then the maximum absorbance peak of the SPR band was measured in each spectrum for the stated days.

### Study of NPs synthesis at low temperature

An assay testing the capability of synthesis of AgNPs was performed using the same procedure as in 2.5 with the strain 2. The strain was grown at 6°C to produce biomass and the biosynthesis of AgNPs was carried out at the same temperature (6°C). Absorbance was measured until day 3. Dry weight of the mycelium was measured.

### Characterization of AgNPs

The AgNPs were characterized using a scanning electron microscope (SEM) model Zeiss SUPRA TM 40 (Oberkochen, Germany) in the secondary electron mode, and with a FEI-Talos F200X G2 FEG Scanning Transmission Electron Microscope in STEM mode combined with a high angle annular dark field (HAADF) detector, 0.16 nm of resolution. The detector allows registering the transmitted beam electrons that have been scattered by the sample through a relatively high angle. Energy-dispersive X-ray spectroscopy (EDS), which is an analytical technique used for the element analysis or chemical characterization of a sample, was conducted on our sample. Taking into account the similarities between de SPR obtained, NPs characterizations were conducted on strain 2.

### Enzymatic reactions

In order to detect the presence of oxidase enzymes, reactions were performed using gallic and tannic acid agar media, and tyrosine to detect the presence of tyrosinase (Stalpers 1978). For the detection of Lignin peroxidase activity (LiP), reactions were assayed in culture media with Azure B (Levin et al. 2005). Additionally, a decoloration test of malachite green was performed to detect the activity of the manganese peroxidase (Levin et al. 2005). The relative intensity of the reaction (+/-), was recorded one week after incubation in the dark at 24 C. Assays were performed for all strains under study.

## RESULTS

### Growth curve of *Tulasnella albida*

A growth curve of dry weight as a function of time was made for each *Tulasnella albida* strain (Fig. 1). All strains showed an initial lag growth phase between days 2 and 4, and an exponential growth phase after that. In the case of strain 1 and strain 2, the deceleration phase seems to start at day 8. The deceleration phase is not observed for strain 3 and strain 4 during the time of the incubation.

**Fig. 1.**
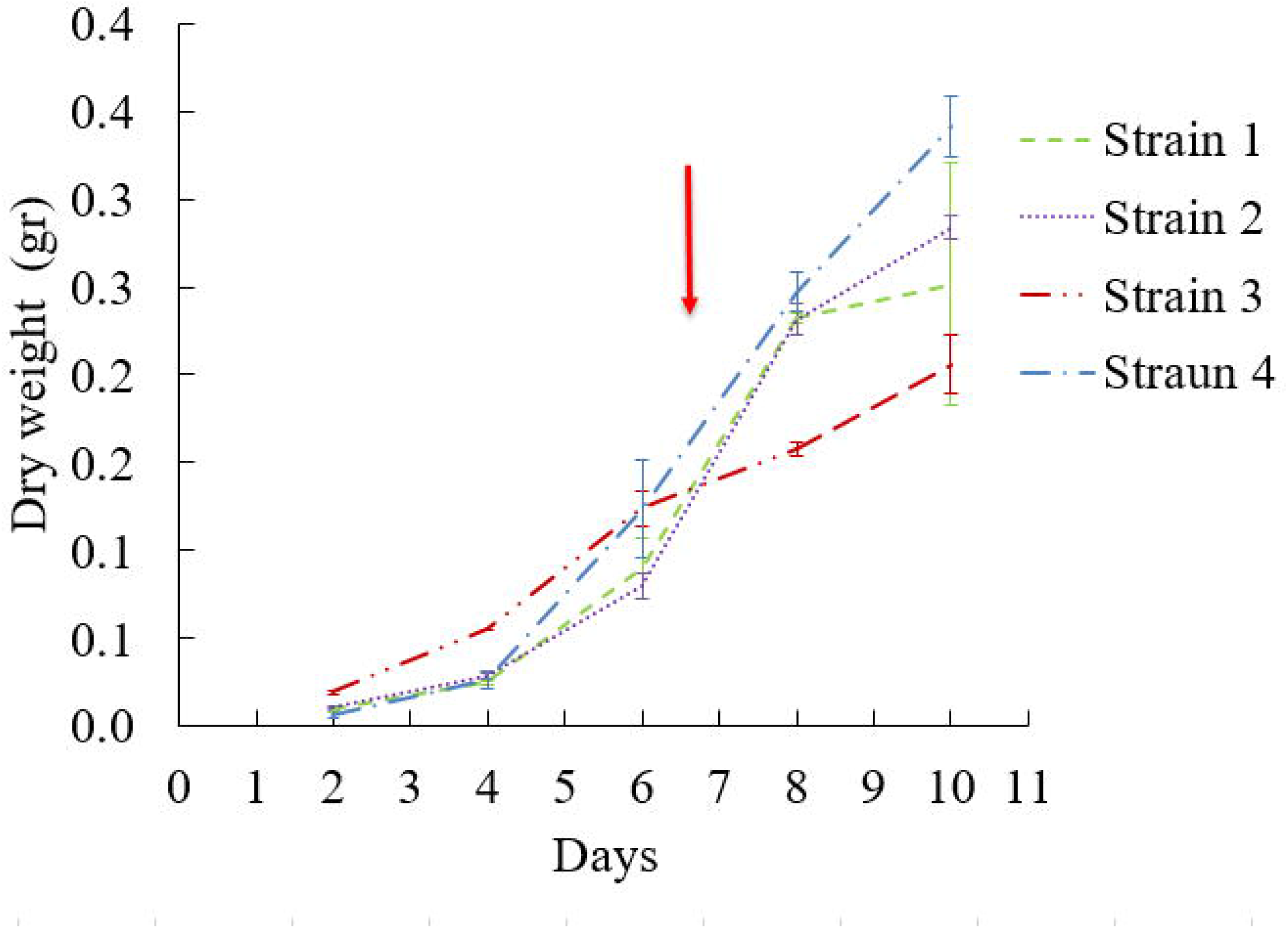
Dry weight growth curve of *T. albida* strains. The arrow marks the moment when the mycelium is harvested for the synthesis

### Biosynthesis of AgNPs

Fig. 2b shows the absorption spectra corresponding to the analyzed strains of *T. albida* on day 7. The color intensity of the AgNPs obtained persisted even after one year, which indicated that the particles were stable in the solution. All spectra show the typical Ag plasmon resonance in the range 400 and 430 nm. A second maximum is observed about 210 nm together with a smaller absorption band in the 230 – 300 nm range. The wavelengths of these secondary maxima agree with the overlapping of silver interband transitions (Shi et al. 2012) and the presence of nanocluster (particles made up of subunits of atoms of Ag) (Ledo-Suárez et al. 2007), and/or could represent a peak of absorbance due to amide band and to tryptophan and tyrosine residue present in the protein molecule (Roy 2017).

**Fig. 2.**
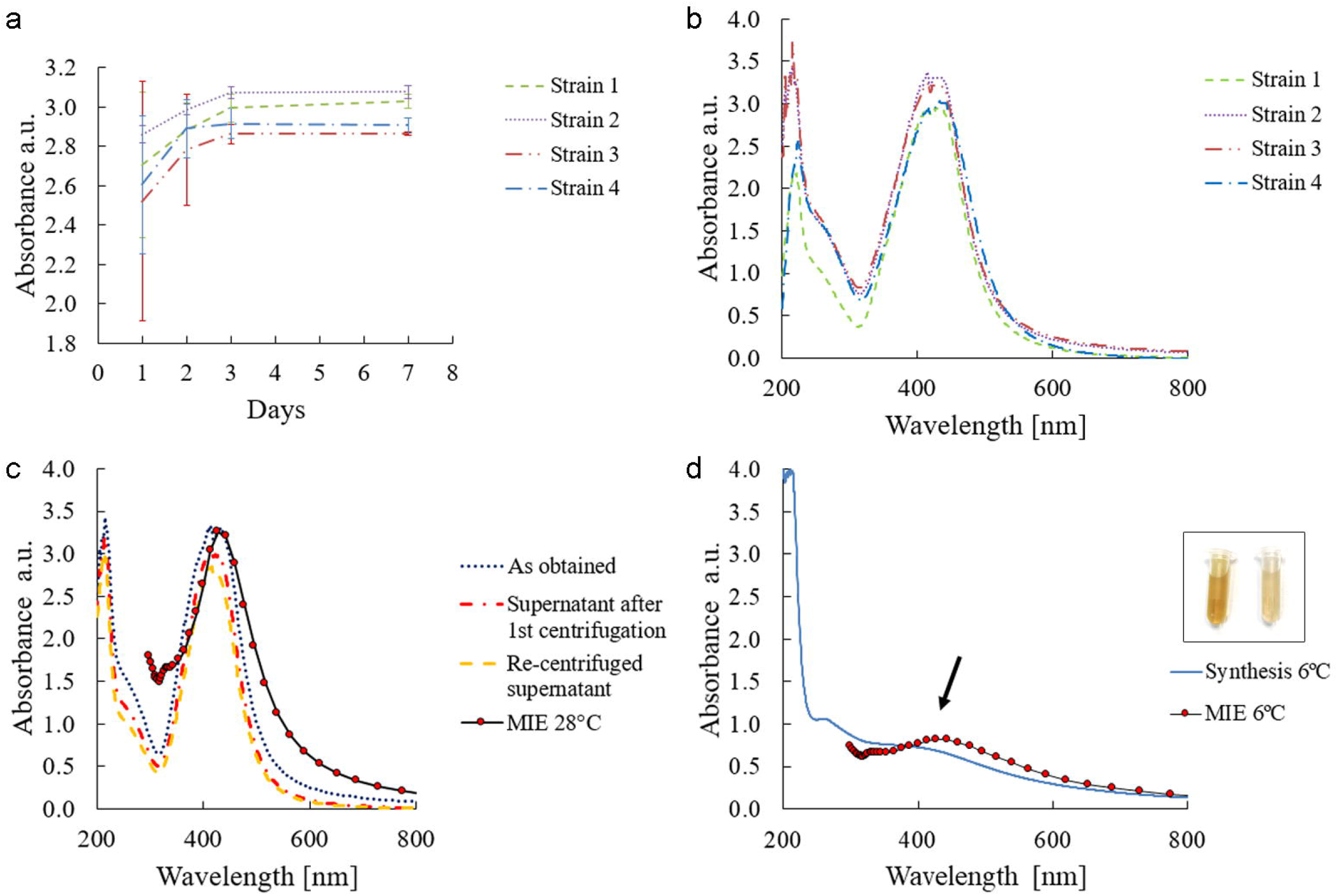
**a**. Maximum absorbance of each *T. albida* strain over time; **b**. Absorption spectra of four different strains of *Tulasnella albida*. Typical Ag plasmon resonance is about 400 and 430 nm together with a second important maximum at 210 nm and a smaller absorption band in the 230 – 300 nm range; **c**. Absorption spectra of the as-obtained colloid (strain 2), supernatant after first centrifugation and re-centrifuged supernatant and theoretical calculation of absorption spectra with 0.9 nm inner radius and 1.05 nm outer radius; **d**. Absorption spectra of the assay at 6°C using the strain 2 recorded on day 3. Typical Ag plasmon resonance at about 400 and 430 nm is emerging (see arrow) and theoretical calculation of absorption spectrum for *Ag-Ag*_*2*_*O* NPs with 0.4 nm inner radius and 0.46 nm outer radius; inset in (d) Nanoparticles obtained at 28°C (left) and 6°C (right)

A mechanical centrifugation was used to explore the possibility of species separation for use in several applications Fig. 2c shows the three spectra of the centrifuged colloids from the strain 2. After centrifugation of the obtained colloid, the supernatant was separated and re centrifuged. This procedure separates the smaller NPs, which are then dominant in the colloid. This fact is observed in the narrowing in the absorption spectrum (Fig. 2c).

Fig. 2c also shows a calculated optical extinction spectrum of a core-shell *Ag-Ag*_*2*_*O* NPs using Mie theory. For this case, a core radius *r*_*1*_ = 0.9 nm and an outer radius *r*_*2*_= 1.05 nm were used. It can be seen that the experimental SPR peak position of the supernatant redispersed colloid is reproduced. This agreement supports the fact that NPs are spherical in shape and have an oxide shell structure. If the NPs were not spherical (spheroidal, rods or triangles) there would be other plasmon resonances at larger wavelengths in the near IR (Noguez 2007).

### AgNPs synthesis efficiency

The gradually progressing reaction of AgNPs synthesis has been monitored using UV–vis spectrometry (since the intensity of the plasmon peak is proportional to the concentration of AgNPs produced) recording the maximum absorbance of SPR on days 1, 2, 3 and 7 of incubation (Fig. 2a). All the spectra exhibit an intense peak around 400-430 nm corresponding to the SPR of AgNPs, implying that bioreduction of the AgNO_3_ has taken place in the presence of the fungal filtrate. Strain 2 was observed to present a higher absorbance than the other strains, suggesting a higher production of AgNPs. It can be noticed that, while strain 4 reaches a stable maximum on day 2, strains 1, 2 and 3 attain their maximum on day 3.

### Study of NPs synthesis at low temperature

The assay performed shows that *T. albida* (strain 2) can grow at low temperatures producing a biomass of 32.6 ± 6.7 mg after 7 days. Fig. 2d shows the absorbance of the synthesis of nanoparticles at 6°C at day 3. Around 400 nm a small plasmon band appears. Compared to the absorbance of the nanoparticles synthesized at 28°C (day 3), the height of the plasmon is around 4.4 times lower (3.071/0.706: absorbance at 28°C/absorbance at 6°C), similar results could be obtained with biomass data, with a relation of around 4.8 (0.1557 mg at 28°C/0.0326 mg at 6°C). A colored colloid of AgNPs (6°C) was obtained showing a lighter color compared with the AgNPs (28°C) (Fig. 2d).

In Fig. 2d, a theoretical calculation of the absorption spectrum for a core-shell *Ag-Ag*_*2*_*O* NPs with 0.4 nm inner radius and 0.46 outer radius is shown. This spectrum shows a good agreement with the experimental plasmon resonance of AgNPs synthesized at 6°C.

### Characterization of AgNPs

Taking into consideration the similar results of the absorbance spectrums it was decided to evaluate only the NPs synthesized at 28°C. The SEM images in Fig. 3a and 3b show isolated and aggregates of synthesized AgNPs. The NPs were not in direct contact even within the aggregates, indicating stabilization of the NPs by a capping agent, probably proteins (Ballottin et al. 2016, Guilger-Casagrande et al. 2019) (Fig. 3b and 3c, indicated with arrows).

**Fig. 3.**
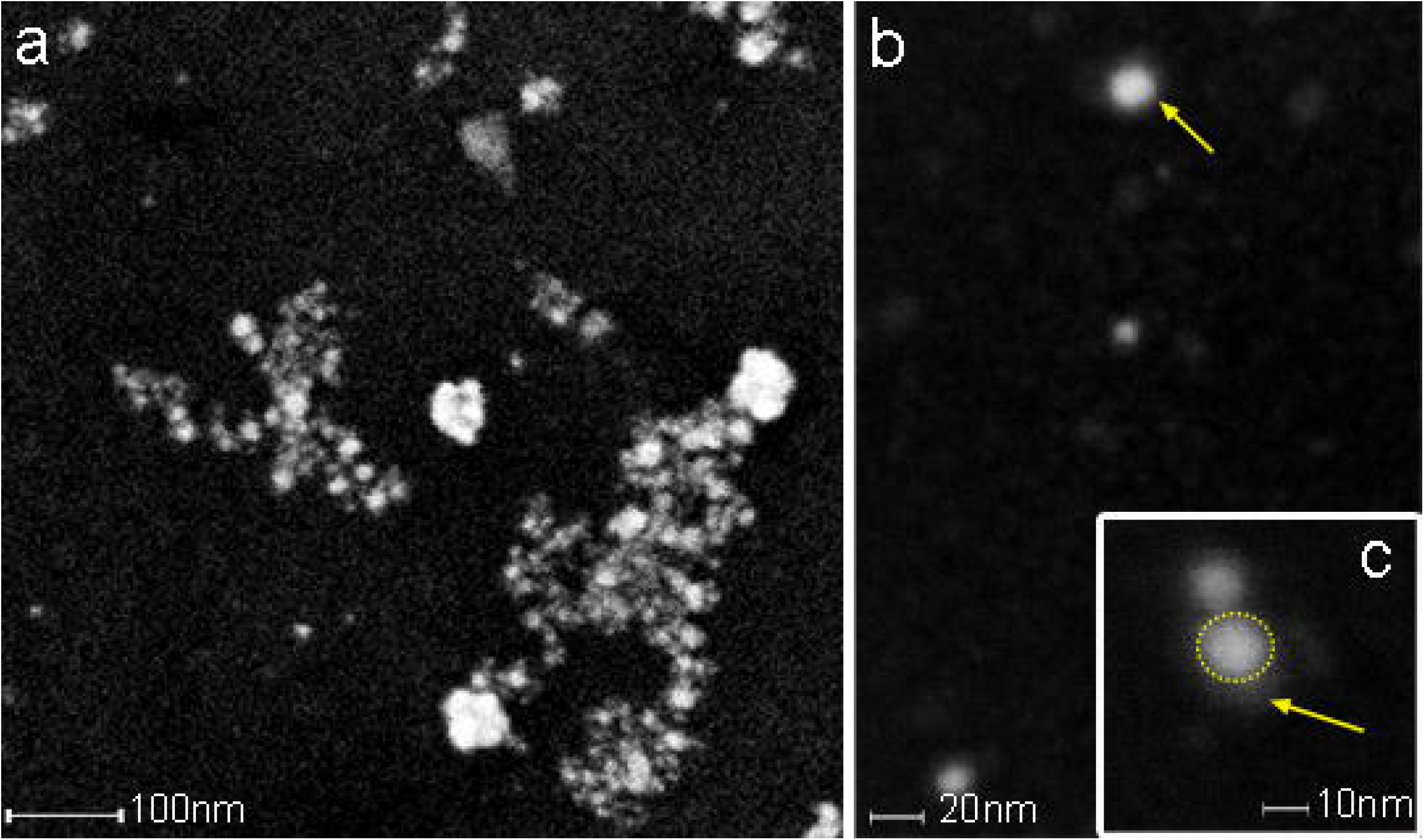
SEM images of colloidal AgNPs synthesized by strain 2 in secondary electron mode (accelerating voltage 3 kV) **a**. Panoramic view of the nanoparticles; **b**. Isolated nanoparticles surrounded by a potential capping structure (yellow arrow); **c**. Detail of a NP (marked by a dotted circle) surrounded by the capping (indicated by the arrow)

Regarding TEM images of synthesized NPs, Fig. 4 shows different magnifications where it is possible to observe AgNPs with spherical shape. Fig. 4a is a panoramic view of the AgNPs. Fig. 4b shows another region with a higher resolution (20 nm). Fig. 4c shows a detail of smaller nanoparticles. Fig. 4d corresponds to the size histogram, showing a size distribution with a maximum at about 2 – 3 nm diam. The most common sources of image contrast are particle mass and crystallinity. Since heavier atoms or massive NPs scatter electrons more intensely than lighter atoms, in dark field mode these regions are brighter.

**Fig. 4.**
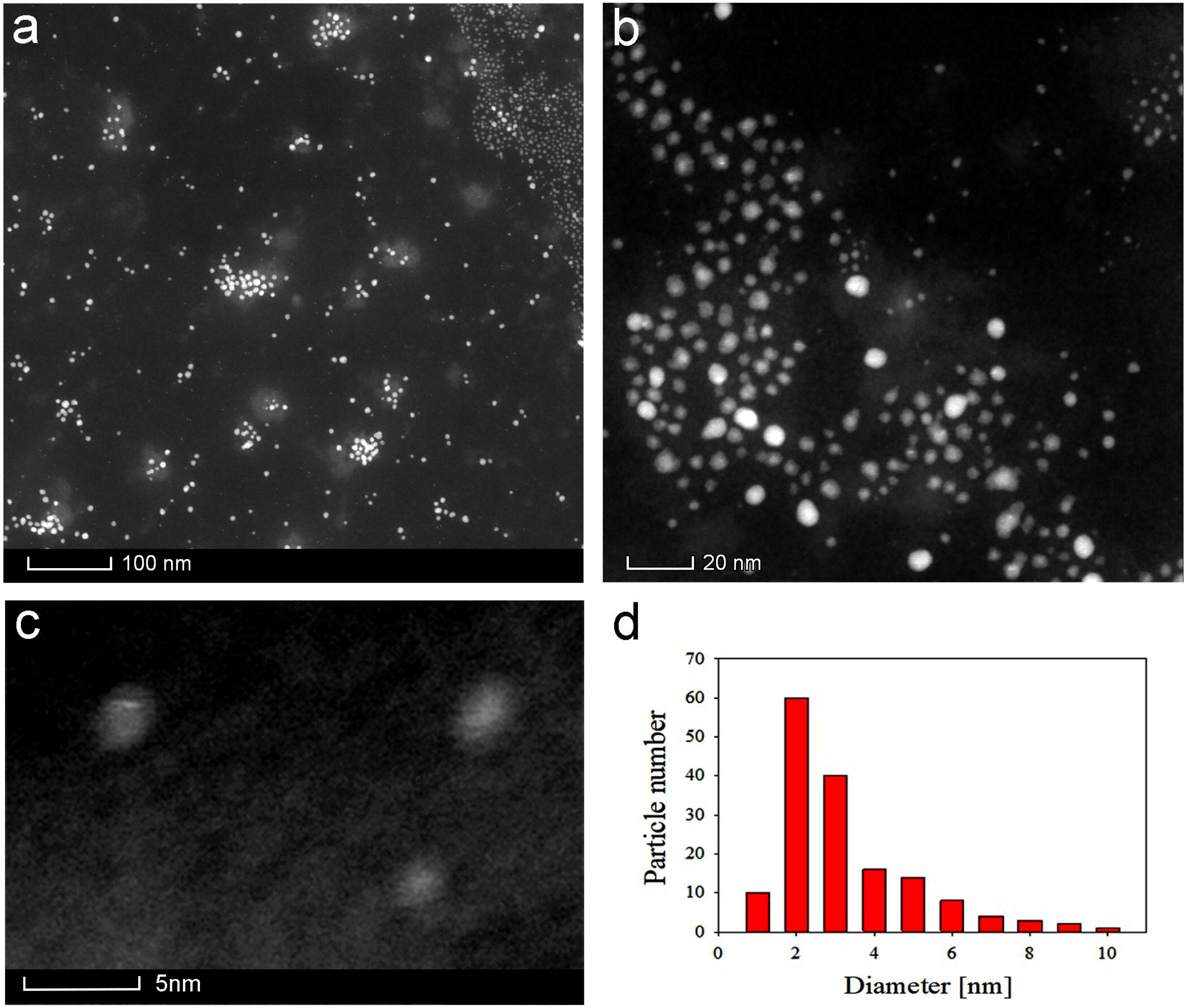
STEM HAADF images of colloidal AgNPs produced by *T. albida* strain 2 **a**. Panoramic view; **b**. Another region in more detail showing spherical NPs; **c**. Detail of smaller NPs; **d**. Size histogram of synthesized NPs

Fig. 5a shows isolated colloidal AgNPs, surrounded by a whitish cloud. Fig. 5b is an EDS from Fig. 5a showing the metal Ag composition of NPs in green dots. It is interesting to notice that the capping surrounding the NPs also yields Ag EDS signals. Green dots are also observed in the background of the image indicating the presence of small amounts of silver. Fig. 5c shows the compositional distribution for selected EDS area at the NPs site of Fig. 5a. The three maxima corresponding to Ag are clearly observed. Fig. 5d shows the compositional distribution for selected EDS area for background signal. Ag maxima are not present in this background. The C and Cu peaks in both panels 5c and 5d arise from the interaction of the electron beam with the sample holder, which is a grid formed by these materials. The O peak that appears in both panels corresponds to the oxidation of metals. The peak in panel 5d is due to oxidation of the copper grid, while the larger O signal in panel 5c corresponds to the sum of oxidation of Cu and Ag.

**Fig. 5.**
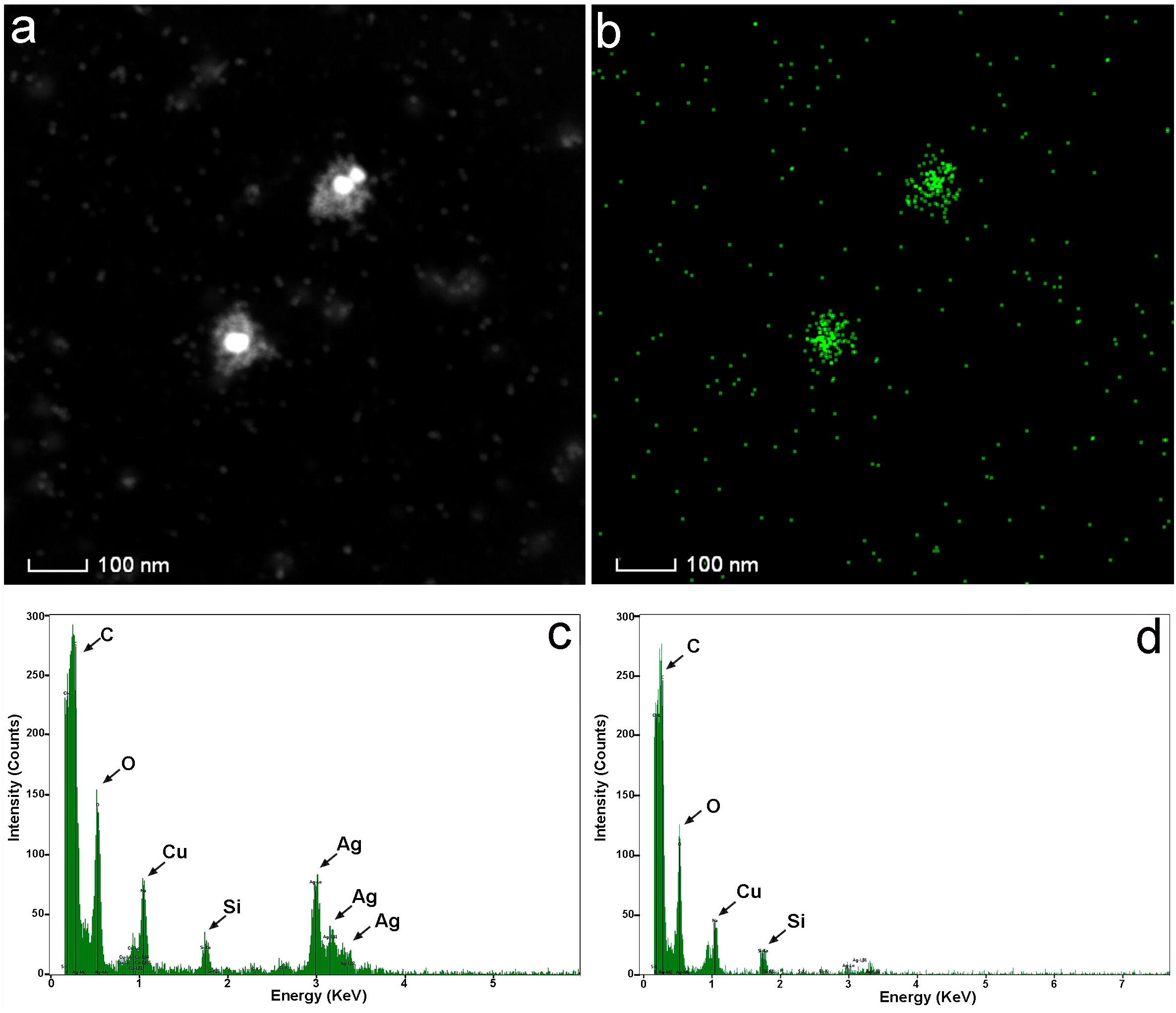
**a**. Dark field images of isolated colloidal AgNPs synthesized by strain 2; **b**. EDS of panel (a); **c**. Compositional distribution for selected area EDS at NPs of panel (a); **d**. Idem (c) for background of the NPs.

Additionally, small amounts of oxygen and carbon could be attributed to the organic layer forming the capping on the synthesized Ag NPs (Jain et al. 2011). Finally, the Si peak corresponds to the signal from the equipment’s silicon detector.

### Enzymatic reactions

Enzymatic reactions were negative for all media and strains evaluated, except for malachite green that resulted in a halo of mild discoloration in all the strains evaluated (data not shown).

## DISCUSSION

In our knowledge, this is the first report of green synthesis performed evaluating psychrotolerant Antarctic filamentous fungi, and the first report regarding *Tulasnella* genus. The study carried out here showed the capacity of these organisms to synthesize AgNPs at suboptimal temperatures both for growth and AgNPs formation process, providing useful data to enhance flexibility of industrial processes.

### Biosynthesis of AgNPs

The data obtained showed that on day 7, when the mycelium is harvested, all *T. albida* strains are in the exponential phase of growth and are capable of producing the metabolites necessary for the synthesis of nanoparticles. The four strains here evaluated showed similar ability to synthesize AgNPs, with strain 2 having the highest efficiency as it obtained higher absorbance, suggesting that the ability to synthesize AgNPs is strain-dependent. Similar results were obtained by Duran et al. (Durán et al. 2005). These authors showed that strains of *Fusarium oxysporum* presented differences in the efficiency of the process, apparently related to the reductase and/or to the quinone generation.

The characteristic plasmon band of AgNPs around 400-430nm of synthesized AgNPs could be seen in Fig. 2b, confirming the presence of these nanoparticles and reaching a stable population after 3 days. No previous reports are known about the use of *T. albida* for the synthesis of metallic NPs.

In this work capped spherical AgNPs (2-3nm approx.) were synthesized using pH 9 and 0.5mM concentration of AgNO_3_ at 28°C. The use of fungi to synthesize metallic NPs is being studied as an effective alternative to the chemical or physical synthesis.

The obtained results here showed that *T. albida* have the ability to synthesize AgNPs at 6°C with biomass (only 32 mg approx.) generated at the same temperature (6°C), showing that strains recuperated from Antarctica could be used in processes of synthesis at suboptimal temperatures. In our knowledge this is the first report of myco synthesis of metal NPs using antarctic fungi studying the synthesis of AgNPs at temperatures as low as 6°C. Analyzing the data resulting from this work, it was observed that the mass and plasmon height relationship between both temperatures were similar (4.4 and 4.8), which could suggest that the synthesis of nanoparticles would be more related to biomass rather than temperature.

Several microorganisms have been used in the synthesis of AgNPs, however, a few reports can be found using psychrotolerant bacteria and fungi (Purcarea et al. 2019). Among fungi *Cryptococcus laurentii* (Ortega et al. 2015) and *Yarrowia lipolítica* (Apte et al. 2013) have been evaluated. On the other hand regarding the study of the synthesis at lower temperatures by microorganisms a few studies were performed. Mageswari et al. (2015) reported that *Pseudomonas mandelii* SR1 synthesized AgNPs with an average diameter of 1.9–10 nm, at 12°C and Javani et al. (2015) reported the synthesis at 4°C of AgNPs using four bacterial strain from Antarctica. Both investigations registered differences with the synthesis at higher temperatures. Meanwhile Mageswari et al. (2015) reported that AgNPs synthesized at lower temperatures (12 and 22°C) are more stable compared with higher temperatures (32 and 42°C). And also, higher temperatures showed distorted plasmon peaks indicating high non-uniformity in the size range of AgNPs and a shift of the UV–visible spectra peak maxima from 410 to 440 nm which was an indication of larger-sized NP production. Javani et al. (2015) found that the average size of NPs depends on the bacterial species and the temperature in an unobvious way, with a trend to smallest size and less dispersivity at lower temperatures. Among filamentous fungi Mohamed Fayaz et al. (2009) and Ahluwalia et al. (2014) registered synthesis at 10°C using *Trichoderma viride* grown at 27°C. The authors reported an increase of size mediated by the temperatures of the synthesis, and the plasmon obtained shift towards higher wavelengths. In this work we did not detect changes in the plasmon during the synthesis at 6°C, however the theoretical model suggests a decrease of the NPs’ size, similar results to the reported by Javani et al. (2015). Our findings indicate that psychrophilic organisms could be an interesting alternative in the synthesis of NPs due to their ability to grow at temperatures below 20°C, to mediate the synthesis at lower temperatures, and to develop studies using the temperature of synthesis as a tool for size control in NPs synthesis.

The second peak observed about 210 nm together with a smaller absorption band in the 230 – 300 nm range, could respond to the presence of Ag nanocluster. Santillán et al. (2020) described bands in the range 200 nm which corresponded to a superposition of silver interband transitions and the presence of nanoclusters. On the other hand, Bhangale et al. (2019) analyzed the synthesis of AgNPs using the fungus *Penicillium* species. They explained the presence of a peak of absorbance at 237 nm due to an amide band and 277 nm to tryptophan and tyrosine residue present in the protein molecule. The presence of the peak at 210-300 nm in this work, even if the spectrum was obtained using the fungal supernatant as a base line, subtracting the effect of the absorption of the amino acids present in the fungal filtrate, appears to support the presence of nanocluster as the better explanation.

Regarding the effect of the centrifugation, it can be observed that as centrifugation steps proceed, the plasmon band decreases in intensity while the UV absorption band remains approximately constant (420-430nm). This fact could indicate that the number of AgNPs decreases faster than nanoclusters in the centrifugation process.

### Characterization of AgNPs

Since the plasmon bands of all obtained colloids were similar, it can be assumed that the morphology of all the nanoparticles is also similar. NPs obtained using *T. albida* resulted in spherical shape of about 2-3 nm, having a capping material surrounding them (Basu et al. 2018). Similarly to our previous work on the synthesis of AgNPs using *Trametes trogii* (Kobashigawa et al. 2019), *T. albida* shows a maximum efficiency of synthesis of NPs on day 3. However, the images obtained from EDS showed that the area occupied by capping has small clusters of Ag, suggesting that the capping could have a role as a reducing agent. In the synthesis of AgNPs using *Lenzites betulina*, Sytu and Camacho (2018) suggest that the capping of nanoparticles is composed of protein and polypeptide molecules. Functional groups such as carboxyl, hydroxyl, and amide groups of the proteinaceous capping molecules could participate in the reduction and stabilization process of AgNPs. The results of the present work are in agreement with the hypothesis that the reduction of Ag could continue at the expense of the capping macromolecules.

### Enzymatic reactions

The exact mechanism of nanoparticle synthesis using fungal supernatants is not yet fully clarified. Some hypotheses have been proposed relating the enzyme production with the fungal synthesis of NPs (Durán et al. 2014, Vetchinkina et al. 2017). Although malachite green test was positive, all other qualitative tests carried out with *T. albida* in this work were negative, so it seems that this fungus does not produce large amounts of enzyme oxidases, peroxidases and tyrosinase (Arora & Sandhu 1984). In this case, *T. albida* ligninolytic enzymes do not seem to be present, having no role in the process of the synthesis of NPs.

The role of NADH dependent nitrate reductase has been already demonstrated (Durán et al. 2005). Regarding *Tulasnella*, it has been reported that *Tulasnella calospora* does not produce this enzyme (Forchi et al. 2017). So, it is possible that *Tulasnella albida* does not produce this enzyme either. Other authors propose the role of polysaccharides and other polypeptides as mediators of the synthesis (Wanarska & Maliszewska2019). Detailed studies evaluating the possible mechanism involved in the synthesis of AgNPs are necessary to elucidate the role of components during the process of synthesis.

## CONCLUSION

The present work provides evidence that Antarctic strains of *Tulasnella albida* are capable of synthesizing AgNPs with a high efficiency at 28°C, and demonstrates that this biosynthesis process occurs at temperatures as low as 6°C. The NPs obtained have spherical shape with sizes between 1 nm and 10 nm. Additionally, our findings suggest the presence of nanoclusters of Ag and its locations indicate that the capping acts as a reducing agent. On the other hand, the mechanism of NPs synthesis remains unknown, however the characterization performed here and the information reported by other authors appears to indicate that neither ligninolytic enzymes nor NADH dependent nitrate reductase have a relevant role in the synthesis of AgNPs in *Tulasnella albida*.

Cold-adapted microorganisms have a great potential in biotechnological application, offering numerous advantages: growth capacity, enzymatic activities and catalytic efficiencies at low-temperature range, preventing the risk of microbial contamination and even in some cases energy saving. These capabilities grant flexibility in industrialization processes, allowing development outside the optimal conditions. This work presents new findings that contribute to the knowledge of extremophilic organisms and their potential use in nanotechnology.

## ACKNOWLEDGMENTS

We acknowledge Dr. A. Caneiro from Y-TEC S.A. Argentina for the use of TEM FEI TALOS F200X, as well as for his commitment and dedication.

## AUTHORS CONTRIBUTION

Kobashigawa, Carmarán and Sacaffardi conceived and planned the experiments. Kobashigawa carried out the synthesis experiments. Schinca and Scaffardi performed the theoretical calculations. Gaiser and Robles performed the enzymatic assays. All authors contributed to the interpretation of the results, provided critical feedback and helped shape the research, analysis, and manuscript.

## Consent for publication

All authors consent to the publication of the manuscript.

